# Genomic signatures of adaptation across a landscape of crickets exposed to an introduced parasitoid

**DOI:** 10.1101/2025.08.30.673289

**Authors:** Jack G. Rayner, Leeban H. Yusuf, Renjie Zhang, Shangzhe Zhang, Oscar E. Gaggiotti, Nathan W. Bailey

**Affiliations:** Centre for Biological Diversity, University of St Andrews; St Andrews, United Kingdom

## Abstract

Novel species interactions provide an opportunity to assess the earliest stages of genetic adaptation in wild populations. We explored the genomic and geographic landscape of adaptation in small, fragmented Hawaiian cricket populations, which are parasitised by larvae of an introduced fly that targets singing males. Multiple protective male-silencing cricket morphs have recently spread despite song’s role in mate attraction. Combining population genomics and field selection experiments, we found sharp declines in the crickets’ effective population size after the fly’s introduction. Nevertheless, geographical variation in selection influences the distribution of male-silencing alleles, consistent with local adaptation. Populations showing the strongest evidence of recovery retain large numbers of singing-capable males, suggesting that initially adaptive responses to the fly created an evolutionary trap via loss of sexual signalling.

## Introduction

Humans impose selection on wild populations in a multitude of ways, but one of the most abrupt is via influencing species dispersal (*1*). From the perspective of evolutionary biology, this offers the otherwise rare opportunity to study populations’ responses to novel selection pressures, e.g., following the introduction of a new predator. In extreme cases, this can foster the emergence of novel coevolutionary interactions, such as those between host and parasite, which can have profound evolutionary consequences (*2*). Understanding the extent to which wild populations are capable of rapidly responding to abrupt changes in selection, how they do so, and the circumstances that follow are key aims in contemporary evolutionary biology. Studies of novel interactions that combine knowledge of the timing of an introduced selection pressure, geographical variation in its strength, and fitness consequences of associated phenotypic variation provide a powerful opportunity to assess genetic adaptation in the earliest stages (*3–5*).

Hawaiian populations of Oceanic field crickets, *Teleogryllus oceanicus*, are locked in a coevolutionary interaction with a deadly eavesdropping parasitoid fly, *Ormia ochracea*. These species are only known to co-occur in Hawaii, to which O. ochracea were recently introduced (*6, 7*). *T. oceanicus* is the primary host for *O. ochracea* in Hawaii; although the fly is also attracted to the songs of other co-occurring cricket species, these species are either infrequently observed (e.g., *Gryllus bimaculatus*) or offer less viable hosts for larval development (e.g., *G. sigillatus*) (*8*). Within the last 25 years, multiple mutant wing morphs have emerged and spread in Hawaiian *T. oceanicus* populations, each of which strongly reduce the amplitude of males’ song. This protects males from being located by gravid female *O. ochracea* (*9–14*), which use cricket song as a phonotactic cue (*15*) and are remarkably accurate in pinpointing the source (*16, 17*), upon which they deposit endoparasitic larvae that burrow into the cricket host. The best characterized of these male-silencing cricket morphs, ‘flatwing’, was discovered in 2003 in a population on the island of Kauai, in which it subsequently spread very rapidly, but has since been observed on two more islands (*9, 18*). However, at least four more alternative male-silencing morphologies have now been described in Hawaiian *T. oceanicus*, and these exist in a heterogeneous mixture of abundances and combinations across a mosaic of small, fragmented populations across the Hawaiian archipelago (*12, 14, 19, 20*). Research into the genetic basis of adaptive responses to selection from *O. ochracea* has focused on identifying loci underlying these mutant wing morphologies, repeatedly finding evidence of large-effect loci or QTL regions (*13, 19, 21*). Yet, while adaptive wing morphs have repeatedly emerged and spread in populations across the Hawaiian archipelago, they have not spread to fixation in most locations. This is likely due in part to strong countervailing fitness benefits of song in the context of attracting and courting females (*22*), suggesting the spread of mutant wing morphs depends on variation in fly attack rates. However, the prediction that the spread of male-silencing phenotypes is influenced by local differences in selection from *O. ochracea* has not been tested.

We characterised the genomic landscape of recent, recurrent and ongoing adaptation across multiple cricket populations to evaluate support for behavioural observations indicating mutant wing morphologies spread under selection from the fly (*9, 18, 20*). We surveyed locations across three islands of the Hawaiian archipelago for *T. oceanicus* populations. From each population (N = 11; Fig. 1A), we resequenced genomes of male crickets (N = 300 genomes at ca. 15x depth of coverage), noting whether each expressed common male-silencing phenotypes (Fig. 1B), and recorded the number of *O. ochracea* that were attracted to cricket song playback as a measure of local selection pressure (N = 489 playback trials across 42 nights). We combine these data for a comprehensive study of the genomic landscape of local adaptation in wild cricket populations, illuminating the conditions that have underpinned this remarkable example of apparently rapid and recurrent adaptation, and its evolutionary consequences.

**Fig. 1.**
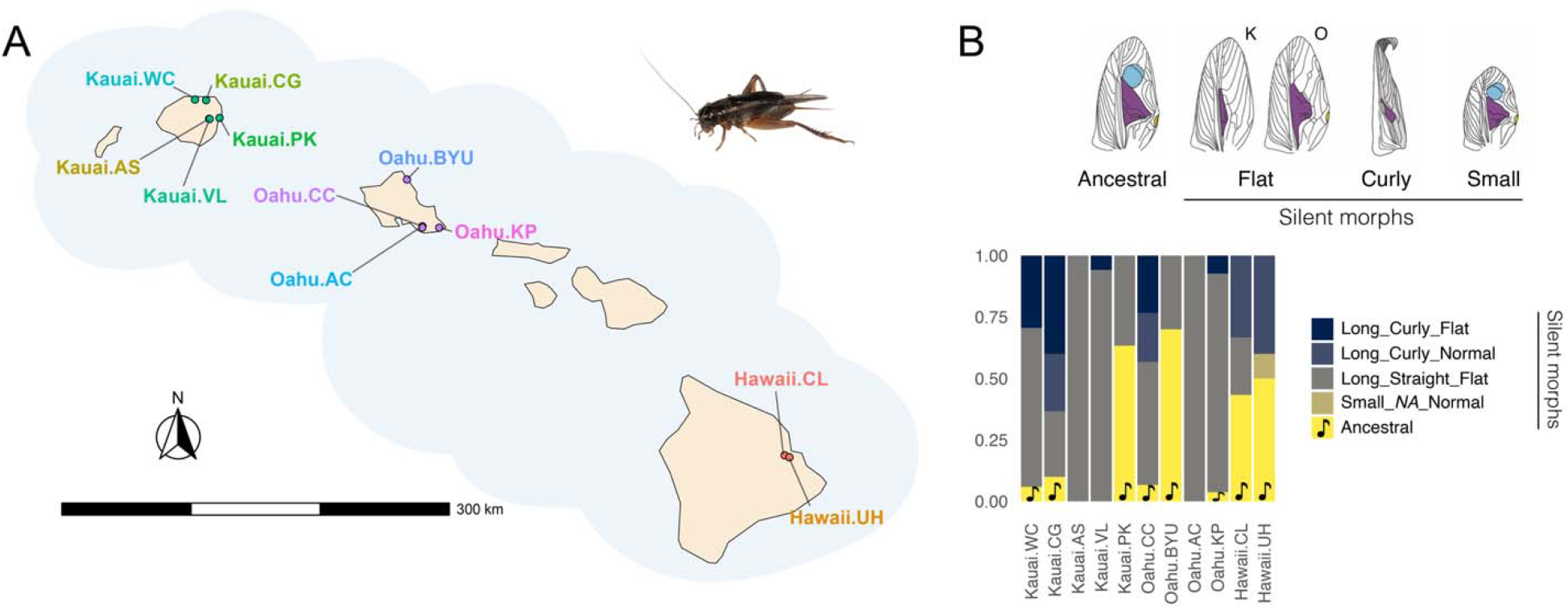
Sampling locations and morphological wing variants in Hawaiian *T. oceanicus*. **A)** Map of Hawaii showing locations from which crickets and flies were sampled. Note that some sampling locations are nearby, but all are separated by urban or geographic barriers and multiple nearby locations show fixed differences in the presence/absence of various wing phenotypes (*19*). Photograph credit: Nathan Bailey. **B)** Proportions of different wing phenotypes (illustrated above) represented in genome-sequencing samples from each patch. Bright yellow bars represent singing-capable males, i.e., those with typical ancestral phenotypes: long wings, straight wing morphology, and normal-wing venation. Small-wing males are assigned ‘NA’ values for wing shape because it is difficult to confidently assess. Line drawings of wing phenotypes are adapted from (*14*). K and O labels above flatwing illustrations represent variants typically found on islands of Kauai and Oahu, respectively.

## Results

### Structural variants dominate genetic and transcriptomic variation

Initial exploratory analyses revealed large structural variants (SVs) dominate genome-wide variation in Hawaiian *T*.*oceanicus*. These SVs were inferred from the presence of discrete linked genomic regions and heterogeneous patterns of population structure across the genome (Fig. 2A, Fig S1) (*23*), leading samples to cluster into discrete karyotypes in chromosome-wide principal component analysis (PCA) (Fig. 2B) (*24*). Based on these criteria, we observed at least one large (> 10Mb) SV on each of 11 of the 14 chromosomes. These SVs were variable in frequency across populations and most appeared absent from ancestral Australian populations (Figs. S2-S3). They include recently described inversions on Chr1 (the X) and Chr2 (*19, 25*), and we did not observe clear differences in read depth in these regions (Fig. S4), suggesting the other SVs are also likely to be inversions. The size of these SVs and their persistence as polymorphisms in small, fragmented cricket populations on multiple islands, presumably reflecting balancing selection, suggests they have important phenotypic effects (*26*). We tested this using existing RNA-sequencing data from neural tissue samples collected from 48 lab-reared crickets of both sexes (*27*). We again found evidence of SVs dominating genetic variation on several chromosomes, and variance partitioning and differential expression analyses revealed very strong local effects on gene expression (Fig. S5).

**Fig. 2.**
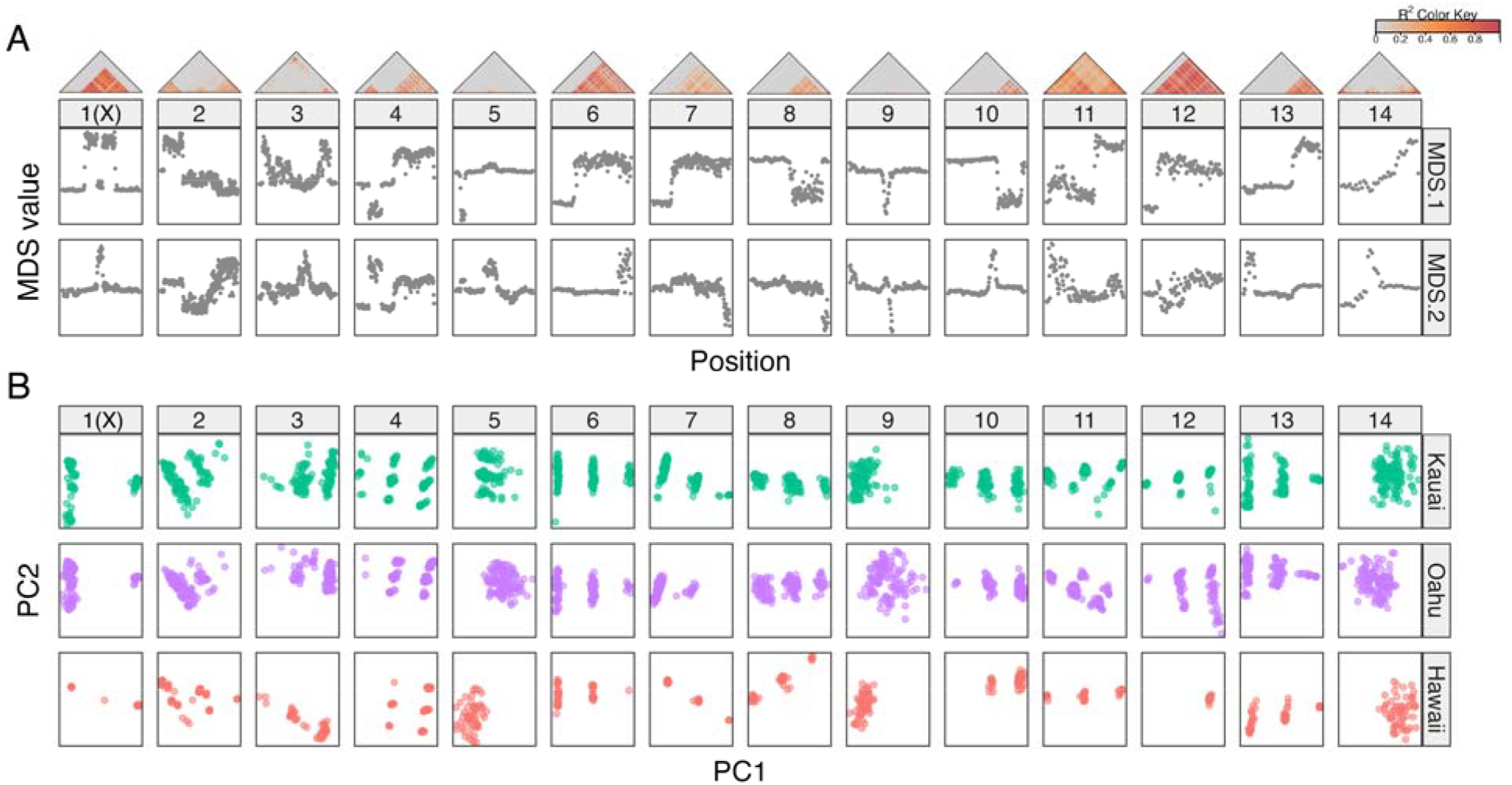
Large, polymorphic structural variants in Hawaiian *T. oceanicus*. **A)** Triangles show observed patterns of linkage across 14 chromosomes. Panels below show MDS values, with discrete tracts of divergence supporting evidence of large structural variants (*23*). **B)** PCA plots by chromosome, separated by island, with discrete clustering of inferred karyotypes into two or three groups illustrating structural variants are typically polymorphic on each island.

### The parasitoid fly is implicated in recent cricket population declines

Because patterns of genomic variation were strongly influenced by SVs, which would likely confound analyses of population structure and demographic inference, we restricted these analyses to linkage-pruned variants from chromosomes 5, 9 and 14, on which we did not observe evidence of large structural variants. Principal component analysis (PCA), approximate maximum-likelihood phylogenetic trees (*28*) and pairwise F_ST_ analyses indicated cricket population structure emerged predominantly at the level of the three islands: Kauai, Oahu, and Hawaii (Fig. 3). There was evidence of finer-scale population structure between sampling locations on Oahu, where *T. oceanicus* appear to have been first introduced to the archipelago (Zhang *et al*., 2021). F-branch statistics (*29, 30*) supported excess allele sharing and thus evidence of gene flow between multiple locations on Kauai, and between locations in Kauai and Hawaii, but not between locations on Oahu (Figs. 3C, S6-7).

**Fig. 3.**
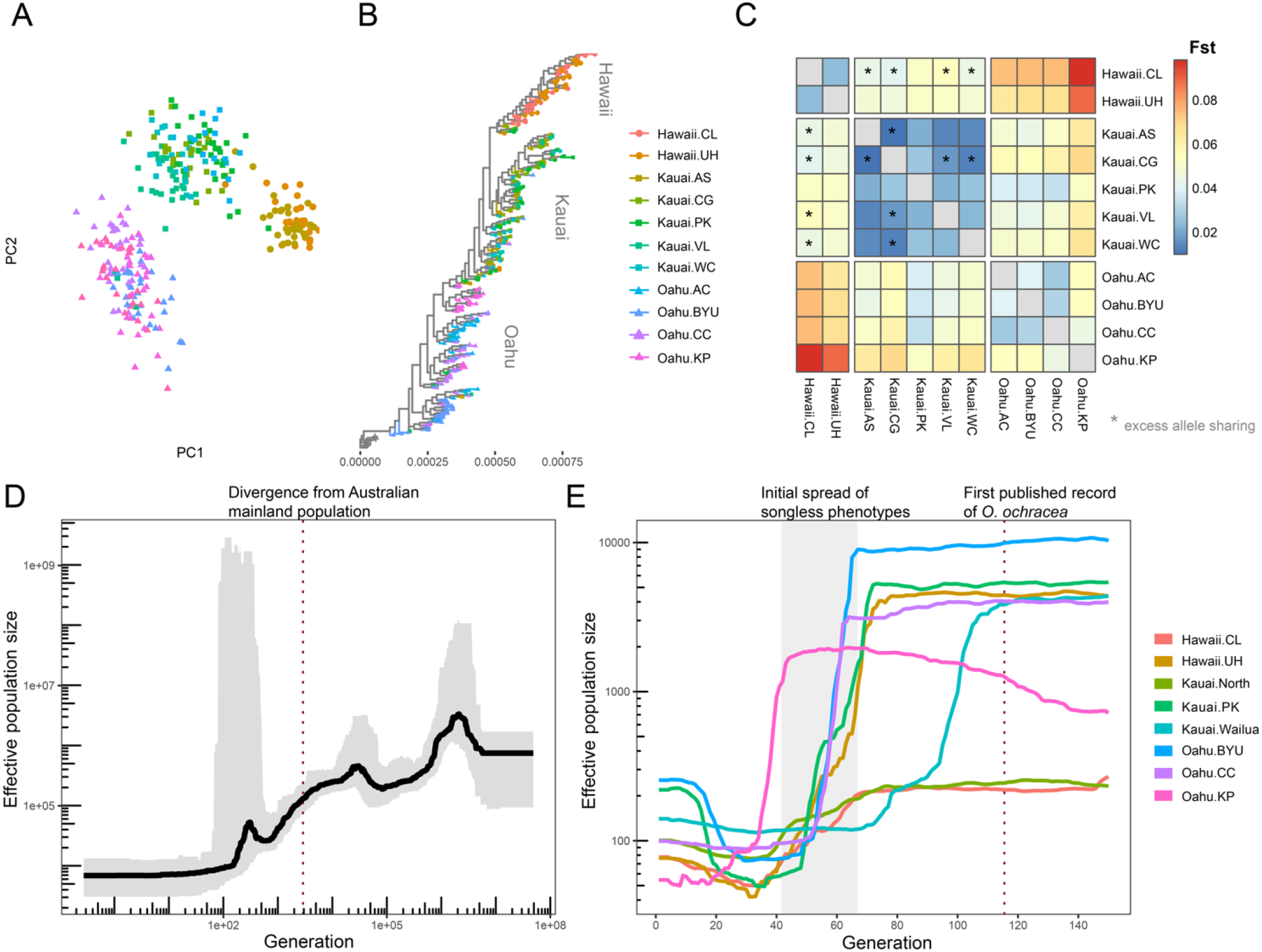
Cricket population structure and changes in effective population size. **A)** PCA of genetic variation across 300 *T. oceanicus* males, with points coloured according to sampling location. **B)** Phylogenetic tree, rooted by the sister species *T. commodus*. **C)** Pairwise mean F_ST_ values across populations. Pairs of locations with excess allele-sharing according to F-branch statistics are identified by asterisks (Fig. S5-6). **D)** Long-term changes in N_e_ inferred from *T. oceanicus* in Oahu. The black line shows the median estimated effective population size. The ribbon illustrates the posterior 95% confidence interval. The dotted line represents the estimated time of divergence between Australian and Hawaiian populations from (*13*). **E)** Short-term changes in N_e_. Lines are coloured according to geographic location of samples. The dotted vertical line indicates the first observation of the parasitoid fly in the Hawaiian archipelago, and the grey rectangle indicates the interval in which the first male-silencing variants were observed to emerge and spread.

*T. oceanicus* appears to have been introduced to Hawaii by Polynesian settlers ca. 800 years ago (*13, 31*), whereas the parasitoid fly was introduced more recently, likely from North America, having been documented in Hawaii since 1989 (*6, 7*). Historical changes in effective population size (N_e_) inferred via Bayesian inference of coalescent times (*32*) indicated a long-term decline in N_e_, likely reflecting repeated bottlenecks associated with divergence from the ancestral Australian mainland population (Fig. 3D). There was a high degree of uncertainty in estimates of effective population size in the interval between 100 and 700 generations before 2022, potentially influenced by fluctuating population dynamics following colonisation of Hawaii, due to factors such as initial population expansion, increasing anthropogenic activity, and the recent introduction of *O. ochracea*.

Assuming an average of 3.5 cricket generations per year (*33*) indicates the fly probably became prevalent within the 150 cricket generations prior to 2022. Linkage-based inference of changes in N_e_ over the preceding 150 generations indicated precipitous recent declines in all populations, fitting expectations following the first documented observation of *O. ochracea* in Hawaii (Fig. 3E). Interestingly, this decline began earliest in the Wailua region of Kauai, in which flatwing was first observed (*9*), suggesting the early emergence of flatwing in this region could have been influenced by unusually strong selection. This is supported by observations from 1993 that crickets at this location were more than twice as likely to be infested by fly larvae compared with those in Oahu or Hawaii (*34, 35*). Declines could also have been influenced by the subsequent spread of songless phenotypes, via reductions in reproductive output due to the reduced frequency of an important signal used in mate attraction. However, two populations (*Kauai*.*PK* and *Oahu*.*BYU*) that retain a high frequency of singing-capable morphs showed particularly severe declines in N_e_ (Fig. 1B cf. Fig. 3E), supporting the primary influence of the parasitoid fly. It is also notable that these two populations also showed the greatest evidence of recent recovery in N_e_, perhaps because large numbers of males retain the ability to attract female mates.

### Selection by parasitoid flies is strongly heterogeneous

In 489 replicate 10-minute trials, cricket song playbacks attracted 415 female flies, whereas an equal number of silent traps attracted just a single fly. While males with mutant wing morphs such as flatwing, curly-wing and small-wing are technically capable of producing sound through stridulation, the amplitude of this ‘song’ is reduced to near-ambient noise levels and prior research has repeatedly shown that fly traps broadcasting their song attract no more flies than those not broadcasting song (*12, 16, 36*). Our observations therefore reiterate that males expressing these mutant wing morphs are very strongly protected from parasitism, resembling findings from dissections of wild-caught Hawaiian *T. oceanicus*, in which fewer than 1% of flatwing males harboured parasitoid larvae contrasting with > 30% of singing-capable males (*9*). Fly attack rates were highest in the first hour following sunset, after which they steadily declined (Fig. 4A), consistent with observations from populations from North American mainland populations (*37*). Across nights, the likelihood of a song playback attracting one or more flies in the two hours after sunset was significantly repeatable with respect to location (linear mixed model with log-transformed response: R = 0.575, 95% CI = [0.178, 0.801], P < 0.001). Nightly fly attack rates were positively associated with increased precipitation (linear regression: F_1,29_ = 12.370, P = 0.001) but did not show a relationship with temperature (F_1,29_ = 0.627, P = 0.400), and differed significantly between locations (location nested within island: F_8,29_ = 9.826, P < 0.001; full model R^2^ = 0.807). Interestingly, at one location with a high proportion of singing male crickets (Kauai.PK), song playback attracted just a single fly in 36 trials across three nights. Conversely, song playback attracted substantial numbers of flies at all three locations in which singing male cricket phenotypes are absent (Fig. 4B), consistent with previous anecdotal observations (*8*).

**Fig. 4.**
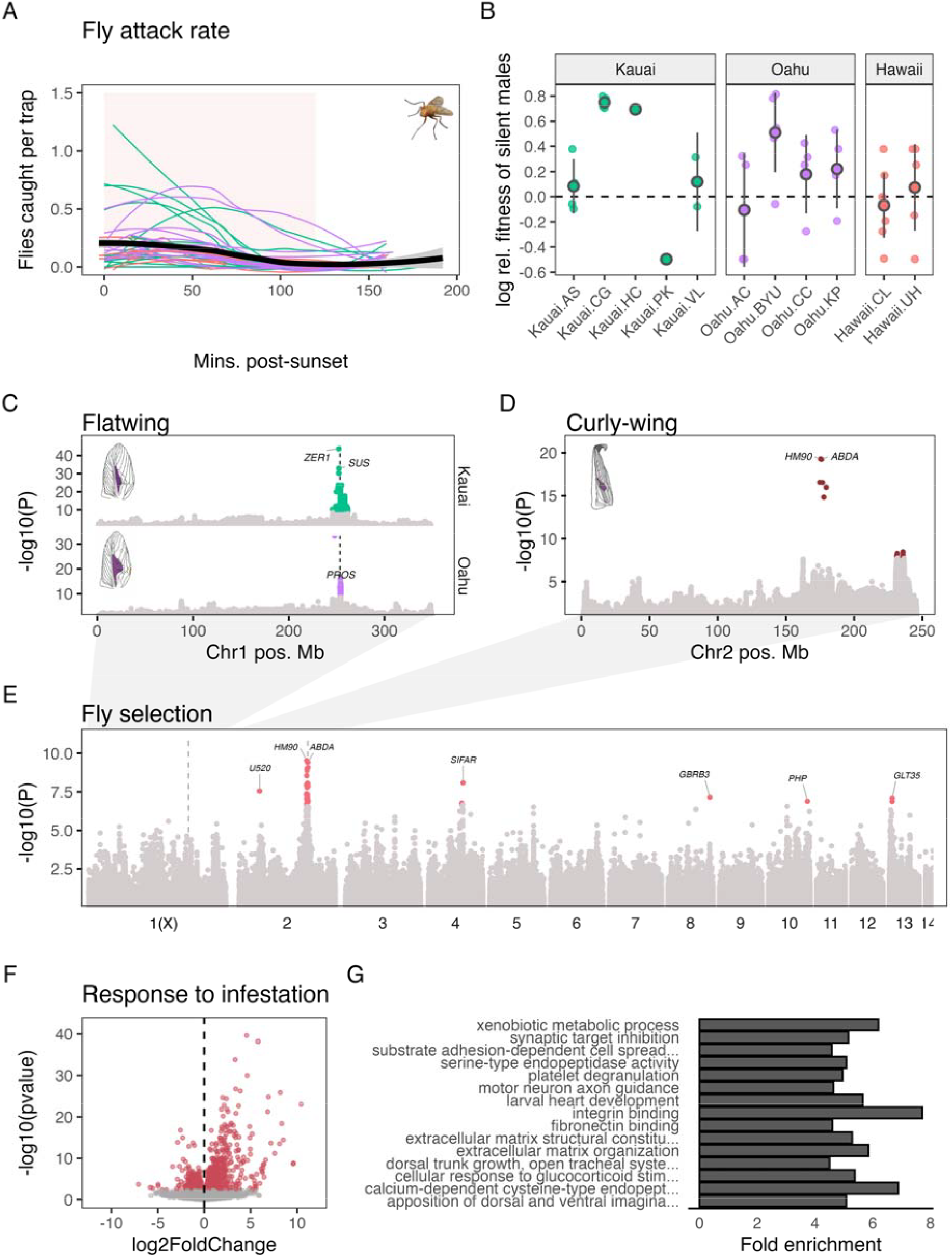
Variants associated with song-loss phenotypes and selection strength. **A)** Numbers of flies caught per 10-minute playback over time. Each coloured line shows loess regression from a single night’s trials, coloured by island, with the black line representing the average number of flies caught across all nights ± 95% confidence intervals. *O. ochracea* photograph adapted from (*12*). **B)** Log-transformed relative fitness of silent males at each site. Small transparent points show estimates per night. Larger points show averages across nights ± 2 standard errors. **C**,**D)** Variants and the statistical strength of their association with flatwing and curly-wing phenotypes. The dashed line in panel C indicates the position of *DSX*. Significant (P_adj._ < 0.05) Fw-associated variants are highlighted in green and purple for Kauai and Oahu flatwing phenotypes, respectively, and Cw-associated variants in dark red. Illustrations of flatwing and curly-wing phenotypes are adapted from (*14*). **E)** Variants and the strength of their association with fly attack rates, with genes near variants with P_adj._ < 0.05 (red) highlighted. Flatwing and curly-wing associated regions are indicated by dashed vertical lines. **F)** Differential gene expression between body tissues of crickets that were uninfested or 4-days post-infestation with fly larvae. Significantly differentially expressed genes are highlighted in red. **G)** The top 15 overrepresented gene ontology categories among differentially expressed genes.

We estimated the average reproductive lifespan of singing-capable males at 16.514 days ± 7.054 SD across nights/locations, based on prior knowledge regarding typical lifespan, the likelihood of infestation having fatal consequences, and the time taken for larvae to develop and kill the host (see Methods and Materials). Relative fitness of reduced-song males in the context of fly parasitism is high but greatly variable across sites, with mean 2.082 ± 0.841 SD. However, silent males are disadvantaged in the context of attracting female mates, and a previous study estimated the relative fitness of silent males in the context of reduced mate attraction at 0.609 (*22*). The net relative fitness of silent males was thus estimated 1.268 ± 0.512 SD, indicating a variable direction of selection across locations (minimum relative fitness = 0.609 at Kauai.PK, maximum = 2.257 at Kauai.CG), and in many cases, across nights (Fig. 4B).

### Selection influences the frequency of a novel male-silencing morph

Testing whether alleles underpinning male-silencing morphs, flatwing and curly-wing, show evidence of selection related to fly attack rates requires knowledge of genomic regions underpinning each morph’s expression. Both have previously been subjected to genome-wide association studies, but with smaller sample sizes (or reduced representation sequencing) and earlier genome assemblies for *T. oceanicus* (*11, 13, 19, 21*), so we reperformed these analyses. A complication is that at least two and perhaps three convergent flatwing phenotypes appear to have arisen independently via independent mutations in the region of the gene encoding doublesex (*11, 13*) on Chr1 (the X chromosome); genetic knockout and knockdown of doublesex is associated with flatwing-like phenotypes in other cricket species (*38*). Prior work has indicated they are stratified by island (*11, 13*), so we identified variants that are consistently flatwing-associated on both Kauai and Oahu (with too few flatwing phenotypes surveyed from Hawaii for a similar analysis), among which it is reasonable to expect signatures of selection across islands. This analysis highlighted 10 Fw-associated variants shared across islands (P_adj._ < 0.05 in Kauai and Oahu), all of which were annotated for the nearest gene *prospero*, adjacent to the gene encoding doublesex, at 254Mb on Chr1 (Fig. 4C). Curly-wing associated loci were located on Chr2, an autosome, near the locations of two homeobox genes at 176Mb (Fig. 4D): *ABDA* and *HM90*.

We compared genetic variation across the 300 cricket genomes from all 11 locations with the nightly probability of a singing male becoming fatally infested using a latent factor mixed model (LFMM) (*39*). QQ-plots of observed versus expected p-values indicated a strong overrepresentation of low p-values in the empirical data (Fig. S8). Of 40 significantly (P_adj._ < 0.05) selected variants identified in this analysis, 33 (82.5%) were within 5 Mb of the consistently curly-wing-associated locus at 176 Mb on Chr2, and were annotated for proximity to curly-wing associated genes *ABDA* and *HM90* (Fig. 4E). Interestingly, the flatwing-associated region was not strongly highlighted, perhaps influenced by its multiple genetic architectures, or because it spread much less recently than curly-wing (*9, 18*).

Other selected variants were within or nearby five more genes distributed across the rest of the genome—annotated for similarity with *Drosophila* genes *SIFAR, U520, GBRB3, GLT35*, and *PHP*—which might reflect selection on other traits important in evading or suppressing larval infestation. One of these, *GLT35*, influences tracheal development, with mutations in this gene leading to malformed tracheal tubes in *Drosophila* (*40*). Tracheal development is directly implicated in cricket responses to larval infestation, due to the apparent ability of *O. ochracea* larvae to co-opt hosts’ immune encapsulation responses to construct trachea that they use to avoid suffocating (*63*). To explore further, we interrogated publicly available RNA-sequencing data from body tissue of Hawaiian *T. oceanicus* that were uninfested, or had been infested for four days (*41*). We found 1,099 genes (of 13,374 in the analysis) significantly differed in expression after 4 days’ infestation (Fig. 4F). None were highlighted by our selection analysis above, however, differentially expressed (Padj. < 0.05) genes were overrepresented in gene ontology categories ‘serine-type endopeptidase activity’ and ‘integrin binding’, both of which are associated with immune encapsulation (*42, 43*), and for genes involved in ‘dorsal trunk growth, open tracheal system’ (all P ≤ 3.10e^-5^) (Fig. 4G). The overrepresentation of these genes supports the view that encapsulation and tracheal development play important roles in larval infestation, and could present a potential target for selection. The apparent trade-off between host encapsulation response and larval co-option to construct tracheal airways has led to speculation that selection might act on attenuated encapsulation responses (*44*), but our work demonstrates it could also act on variants influencing tracheal development.

## Discussion

The recent introduction of a parasitoid fly to the Hawaiian archipelago exposed local *T. oceanicus* populations to a novel and extreme selection pressure, offering rare insight into the circumstances of rapid adaptation. The strength of selection is evidenced by field assays of fly attack rate and severe declines in effective population size. We show that this has created variation in the frequency of alleles underpinning the common male-silencing ‘curly-wing’ phenotype, though interestingly not of another male-silencing ‘flatwing’ phenotype that preceded the emergence of curly-wing. This could be because flatwing is fixed in or absent from multiple populations, in which its frequency cannot change without introduction of other alleles via gene flow, but it could also be due to independent flatwing-associated mutations (*13*). However, we also observe evidence of fly selection influencing the frequency of other genetic variants, many of which correspond to traits of interest in the coevolutionary interaction between crickets and the fly parasitoids.

Genetic adaptation to anthropogenic selection pressures can have negative long-term impacts upon wild populations, by drawing them away from previous local fitness optima (*45*). Songless male *T. oceanicus* have greater fitness in the context of the parasitoid fly but are strongly disadvantaged in the context of sexual selection. Our estimates of net selection indicate that silent males are strongly favoured at three of the 11 locations surveyed, but are strongly selected against in one population, Kauai.PK. Across the remainder, the direction of selection varied nightly, though these include three locations at which singing-capable males are absent (*12, 46*). It is noteworthy that Kauai.PK and Oahu.BYU, two populations retaining the largest proportions of singing male phenotype, also showed the strongest evidence of recent recovery in effective population size following severe initial bottlenecks. This is intuitive, because singing-capable males are much more likely to reproduce. We also observed non-negligible fly attack rates at three locations in which no singing-capable male morphs remain, which would be expected to greatly reduce parasite populations (*47*). This is curious, but a recent study showed flies are also attracted to the song of other co-occurring Grylline crickets that offer less viable hosts (*8*), raising the intriguing prospect that *O. ochracea* are able to co-opt the acoustic signals of other species to parasitise silent *T. oceanicus* populations, complicating the coevolutionary interaction. An alternative possibility is that flies are itinerant and were attracted from nearby locations. We have not heard singing males in recent surveys of areas surrounding these sites, but the likely migration asymmetry between flies and crickets represents an important direction for future research, as this is key to predictions of local adaptation (*48*).

Besides male-silencing phenotypes, we detected further evidence of selection on variants within or near genes corresponding to traits that attracted the interest of early behavioural studies investigating adaptive responses to the parasitoid fly (*34, 44, 49*)—in some cases prior to the emergence of silent male morphs. One selected variant was within the gene *SIFAR*, which in *Drosophila* influences variation in wing extension index during male courtship (*50*). Prior work in Hawaiian *T. oceanicus* explored whether singing behaviour differs across populations (*34, 35, 51*), with an early study reporting evidence that singing-capable males in parasitised populations reduced calling effort during the interval of peak fly activity (*34*). More recently, the prediction that selection would act to modulate singing behaviour has been largely overlooked following the spread of adaptive male-silencing phenotypes. We also observed selection on the gene *GLT35*, which influences tracheal development; mutations in this gene lead to malformed tracheal tubes in *Drosophila* (*40*). This, too, relates to prior research in this system. As in other insect hosts of parasitoids, crickets mount an immune encapsulation response against infesting *O. ochracea* larvae (*44*). However, *O. ochracea* larvae appear to co-opt this encapsulation response to construct a tracheal funnel which attaches to the cricket’s abdominal wall, allowing developing larvae to avoid suffocation (63). Our re-analysis of differences in gene expression between uninfested and infested cricket body tissues (*41*) showed infestation-responsive genes are overrepresented not just for functions involved in larval encapsulation, but also tracheal development. The apparent trade-off between host encapsulation response and larval co-option has led to speculation that selection might act on attenuated encapsulation responses (*44*), but it could also feasibly act on genes involved in tracheal development. Whether evidence of selection on genes implicated in singing behaviour and tracheal development reflects adaptive variation in these traits represents an interesting avenue for further research.

We previously reported evidence that the co-occurrence of multiple male-silencing wing morphs could act to impede any one of them from spreading to fixation (*19*). In addition, genomic regions underlying flatwing and curly-wing phenotypes are nearby but not intrinsically linked with large segregating inversions (*19, 25*). We were surprised to find evidence of similarly large segregating structural variants (many if not all of which are likely to be inversions) across nearly all *T. oceanicus* chromosomes, most of which were polymorphic on all three islands, indicating they may be maintained as balanced polymorphisms. These structural variants account for a massive proportion of the genetic variation and gene expression in these populations, suggesting they have substantial phenotypic effects. As well impeding recombination between potentially adaptive loci, selection acting upon these large SVs could be a source of interference that impedes the spread of adaptive variants (*52*). The maintenance of inversions at intermediate frequencies has received considerable recent interest (*53*), but the contribution of different potential sources of balancing selection on these large regions are still not well understood (*54*). A combination of theoretical and empirical studies have implicated various factors influencing balancing selection in large linked regions, including overdominance or heterokaryotype advantage (*55, 56*), epistasis and antagonistic pleiotropy (*57*), and sexual antagonism (*58*). Hawaiian *T. oceanicus* appear remarkable for the number of large potentially balanced structural polymorphisms that are maintained at intermediate frequencies within several populations. Given the strength of selection that is likely to be necessary to maintain inversion polymorphisms in multiple small populations, and their evolutionary consequences, Hawaiian *T. oceanicus* could offer a compelling system to comprehensively test the sources of balancing selection maintaining variation in these regions.

## Materials and Methods

### Selection assays

We performed 489 fly capture trials starting at sunset on 42 nights at 11 patches, and were performed in January-February and June-July in 2022. Each fly capture trial involved eight traps, half of which broadcasted a cricket song model based on average song parameters across several Hawaiian populations of *T. oceanicus* (*59*), and lasted 10 minutes, with song broadcast at 85dB at a distance of 5cm using a speaker (Jam HX-P303BK) attached to an mp3 player (SanDisk SDMX26-008G-E46P). Control traps were identical to those broadcasting song, but without playback. Traps were set out in a 3 x 3 square, with 1.5m between traps and an alternating order of control and song traps, which were swapped each trial. Trials began at sunset, with twelve trials per night (unless interrupted by heavy rain). After trials, captured flies were released 10 metres from the trap, after which we waited for three minutes before resuming trials. We recorded the number of flies attracted to each trap. One site, Kauai.WC, was only assayed on a single night in October 2022. Because fly abundance was repeatable across nights at other sites (see main text), did not appear to differ between surveys performed in early and mid-2022, and because assays at this site reported similar rates of fly attack as a nearby site, Kauai.CG, we retained this site in our analysis.

Repeatability of fly attack rates across nights were estimated using rptR v0.9.22 (*60*) by running linear mixed models with a response variable of the log-adjusted fly attack rate between within 2 hours of sunset, averaged across nights. Data for average daily temperature and precipitation at each location were retrieved using the nasapower R package.

### Estimating expected lifespan

Attracting a single fly has drastic fitness costs, as infestation by *O. ochracea* larvae is very often fatal and gravid females are remarkably accurate in locating the source of cricket song (*16, 17*) even after the cessation of singing behaviour (*61*). To estimate the expected adult lifespan of singing-capable males at different locations, from which to infer the strength of selection against song, we made the following assumptions based on existing data. Assumption 1: we estimated the typical adult lifespan in the wild of reduced-song (e.g., expressing curly-wing or flatwing morphs) males at 29 days, based on longevity records from a wild population of a related field cricket, *Gryllus campestris* (average longevity of 28.9 days) (*62*) and from *T. oceanicus*’ sister species, *T. commodus*, reared under semi-natural conditions (29.07 days) (*63*). Assumption 2: males sing for an average of 7.6 minutes per night in the two hours following sunset, as observed in a prior study (*51*), and begin singing shortly after adult eclosion (singing behaviour was recorded at 0-3 days post-eclosion by (*64*)). Assumption 3: phonotaxis by gravid female *O. ochracea* has fatal consequences for *T. oceanicus* males 61% of the time, based on findings from a related cricket host, *Gryllus texensis*, in which deposited planidia successfully established in, emerged from and killed 61% of males (*65*). Finally, assumption 4: Cessation of reproductive viability occurs at six days post-infestation; Adamo et al. (*66*) found reproductive viability strongly declines after five days infestation, with larvae emerging and killing crickets 7-10 days post-infestation.

Following the assumptions stated above, we calculated the number of flies a singing-capable male cricket attracts per day as *N*_*fly*_ = mean number of flies attracted per 10-minute playback * 0.76, based on prior observations of nightly singing effort (*51*). We then assume:

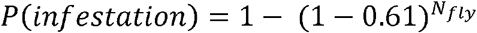

Where *P(infestation*) is the daily probability of becoming fatally infested, given that fly planidia successfully infest and emerge as larvae from 61% of male *G. texensis*, another cricket host (*65*). We further assume that the lifespan of a cricket follows a geometric distribution, with a time lag *L* between infection and loss of reproductive viability (6 days – see above). Under this assumption, a singing male’s expected reproductive lifespan is:

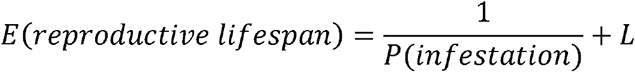

On top of this, we imposed a maximum expected reproductive lifespan of 29 days, the same as that of silent males.

### Sample collection, extraction, sequencing

Tissue samples were collected from wild-caught males at 11 locations between July 2021 and June 2022 by removing a single hindleg. Samples from one population (Oahu.AC) were supplemented with samples collected in October 2018 (N = 15 of 30). This population was therefore not included in analysis of recent changes in effective population size. After tissue collection, animals were released, and crickets missing a hindleg were not sampled to avoid resampling. Legs were stored in 100% ethanol at -80C until DNA extraction using a CTAB-chloroform protocol. Libraries were prepared using NEBNext Ultra II FS Kits with ∼300bp inserts, and sequenced at the University of Liverpool Centre for Genomic Research on an Illumina NovaSeq S4, generating paired-end 150bp reads. Ninety of these sequenced samples were used in a previously published analysis (*19*).

### Variant identification, calling, filtering

Reads were provided by the facility after initial quality and adapter trimming using Cutadapt v1.2.1104 (*67*) and Sickle v1.200, with a minimum window quality score of 20. Further trimming was performed using FastP v.0.24.0 with default parameters. Reads were aligned to the *T. oceanicus* genome v3 (*25*; ENA accession GCA_976985615) using BWA-mem v.0.7.19 with default parameters, following which PCR duplicates were flagged using SAMBAMBA v.1.0.1. We performed variant discovery from aligned reads of 55 samples, 5 per location, selecting samples with high sequencing depth and which represented all morphological variants recorded at the respective location. Variant discovery and calling of samples was performed in Freebayes v1.3.6 (*68*). Variants were initially filtered using vcftools v0.1.16 (*69*) to retain genotype calls with: minimum sequencing depth greater than 5x and less than 60x; minimum genotype quality 20; minimum quality 30; minimum genotype call rate across samples > 0.75; mapping quality >= 30; biallelic variants (N = 75,180,819 variants). For most analyses, variants with minor allele frequency < 0.05 (or < 0.1 in genetic association and selection analyses) were also removed (N = 48,363,493 variants), although not when quantifying nucleotide diversity or Tajima’s D. Tajima’s D, nucleotide diversity and Weir and Cockerham’s F_ST_ were calculated in 50Kb windows with 10Kb step size in vcftools.

### Windowed PCA and linkage analysis

We used the Lostruct R package to run windowed PCA (*23*), often used to identify putative chromosomal inversions, across the genome. We first further filtered the full VCF for SNPs with less than 1% missing genotypes, samples missing genotypes at less than 20% of loci, and retained only one variant per 50Kb (N = 149,673; 294 samples). We converted the VCF to TPED format and implemented PCA in 50-SNP windows across the genome. To visualise linkage across chromosomes, we thinned the set of SNPs using a window of 200Kb, filtered for minor allele frequency > 0.2, then calculated and plotted r^2^ values in LDheatmap (*70*).

### Population structure and demographic inference

Inferences of population structure and demographic history were performed using linkage-pruned (plink: indep-pairwise 50 10 0.1) variants genotyped in >=90% of samples and located on chromosomes 5, 9 and 14 (those not showing evidence of massive structural variants) (N = 73,674 variants). Phylogenetic relationships were explored using FastTree (*28*), and were plotted using the R package ggtree (*71*). F-branch statistics were calculated in Dsuite (*30*).

Repetitive regions identified by (*25*) were excluded before running analyses to estimate historical changes in effective population size. Long-term changes in effective population size were inferred using samples collected from Oahu (to which *T. oceanicus* were first introduced) in 2022. This analysis was performed in Phlash (*32*). To explore more recent shifts in effective population size, we used GONE2, a method based on patterns of linkage disequilibrium (*72*). For this analysis, we pooled geographically nearby populations (Kauai.CG and Kauai.WC; and Kauai.AS and Kauai.VL) in an effort to avoid confounding effects of possible recent gene flow between these locations.

### Genome-wide association and selection tests

Genome-wide association tests for curly-wing and flatwing phenotypes were implemented in GEMMA (*73*), using linear mixed models with a genetic relatedness matrix included as a random effect, and tested using likelihood ratio tests. Tests for loci that differed in frequency in association with fly abundance were implemented using latent factor mixed models that account for unmeasured variation (i.e., hierarchical population structure) implemented in the R package LFMM (*39*) with ridge penalty estimates and a K value (reflecting the number of populations) of 3, which was determined from inspection of PCA screeplots and the number of islands. P-values were calibrated by a genomic inflation factor. Input data were linkage-pruned in 50-variant windows using Plink (*--indep-pairwise 50 10 0*.*1*), variants missing in > 90% of individuals or with minor allele frequency < 0.1 were removed, and missing genotypes were imputed using the *impute* function in the R package LEA (*74*). Gene annotations were retrieved from (*25*).

### Gene expression analyses

Trimmed, paired-end RNA-sequencing data collected from neural tissue of 48 lab-reared *T. oceanicus* of both sexes (*27*), and body tissues from 18 *T. oceanicus* that were or were not infected with *O. ochracea* larvae (*41*), were downloaded from the NCBI Sequence Read Archive. In each case, reads filtered with FastP and default parameters, then aligned to the *T. oceanicus* genome using HISAT2 with default parameters (*75*), after which gene counts were estimated using StringTie (*76*). In addition, for the 48 samples from (*27*), variant calling was performed using bcftools v.1.21, and filtered for genotype quality (> 20), missingness (< 0.25) and minor allele frequency (> 0.05), before running chromosome-level PCA in Plink (*77*). Variance partitioning of gene expression counts was performed using the R package variancePartition v1.34.0 (*78*), and differential expression analyses were performed in DESeq2 v1.44.0 (*79*). Gene ontology overrepresentation tests for biological processes were performed using topGO v2.56.0 (*80*) using gene ontology annotations retrieved from (*25*).

## Acknowledgments

We are grateful to landowners at all locations for permission to collect crickets on their grounds. We thank Susan L. Balenger for contributing tissue samples. We thank Jessie Tanner for providing cricket song playback models used in selection assays, and Robin Tinghitella, Dale Broder and Gabrielle Welsh for advice in the field. This study was supported by funding from the Natural Environment Research Council (NE/T0006191/1) to N.W.B., O.E.G., and J.G.R. We acknowledge computational resources from the James Hutton Institute Bioinformatics HPC (BBSRC grant BB/S019669/1). We thank Tanya Sneddon, Audrey Grant, Megan McGunnigle, and David Forbes for assistance with laboratory work, and Ana Drago Rosa for assistance with fieldwork.

## Funding

Natural Environment Research Council NE/T0006191/1 (NWB, OEG, JGR)

Biotechnology and Biological Sciences Research Council BB/S019669/1

## Author contributions

Conceptualization: JGR, OEG, NWB

Methodology: JGR, OEG, NWB, LHY, SZ

Investigation: JGR, OEG, NWB, LHY, RZ, SZ

Visualization: JGR

Funding acquisition: NWB, OEG, JGR

Project administration: NWB, JGR

Supervision: NWB, OEG

Writing – original draft: JGR

Writing – review & editing: JGR, OEG, NWB, LHY, RZ, SZ

## Data and materials availability

Newly published genome sequencing data are available at the NCBI Sequence Read Archive under accession PRJNA948842. Previously published genome sequencing data from Hawaiian *T. oceanicus* are at the NCBI SRA under accession PRJNA1019311, and for Australian populations of *T. commodus* and *T. oceanicus* are available from the ENA under accession PRJEB39125, previously published RNA-sequencing datasets are available from the NCBI SRA under accession PRJNA344019 (song experiment) and the NCBI Gene Expression Omnibus under accession GSE15139 (infestation experiment). Scripts used in analyses are available at https://github.com/jackgrayner/cricket_popgen. Additional survey data and results files are made available as supplementary material.

