## Supplemental Information for "Genomic signatures of adaptation across a landscape of crickets exposed to an introduced parasitoid"

**Affiliations**

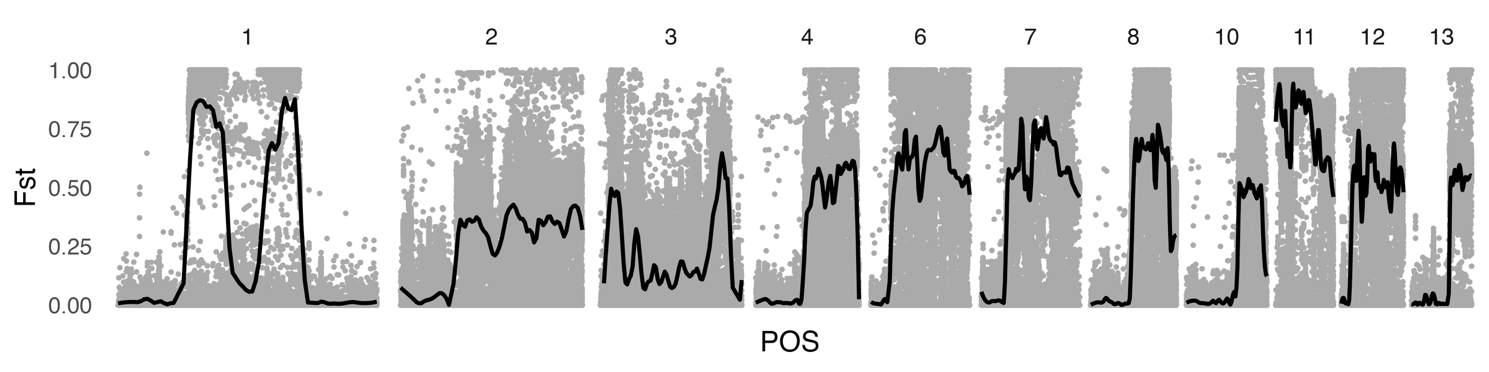

**Fig. S1.** F_ST_ values between alternative homokaryotypes are consistently high in structural variants, with the exception of Chr3, which appears to show a U-shaped pattern with higher divergence near the presumed edges of the variant. F_ST_ values were calculated using just cricket samples from Kauai to avoid confounding effects of genetic divergence between island populations. As above, this does not account for all of the large SVs we observed (e.g., we observed two large SVs on Chr2), but only those that contributed most strongly to variance on PC1. Lines illustrate LOESS trends.

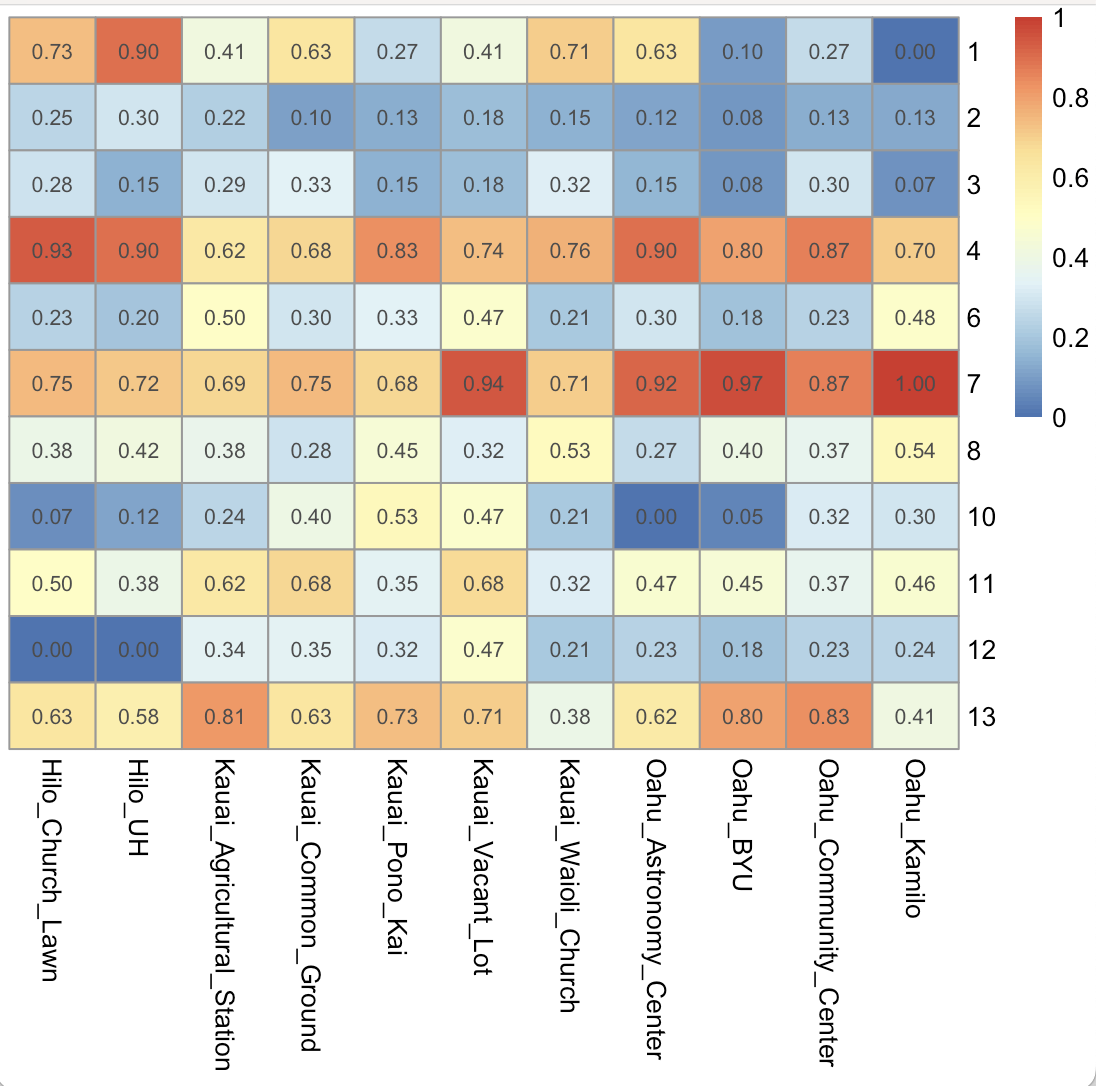

**Fig. S2.** Inferred frequencies of large structural variants inferred from principal component axis 1 (see Fig. 2) on each of 11 chromosomes (rows) in each of 11 locations. Note that this does not account for all of the large SVs we observed (e.g., we observed two large SVs on Chr2), but only those that contribute most strongly to variance on PC1.

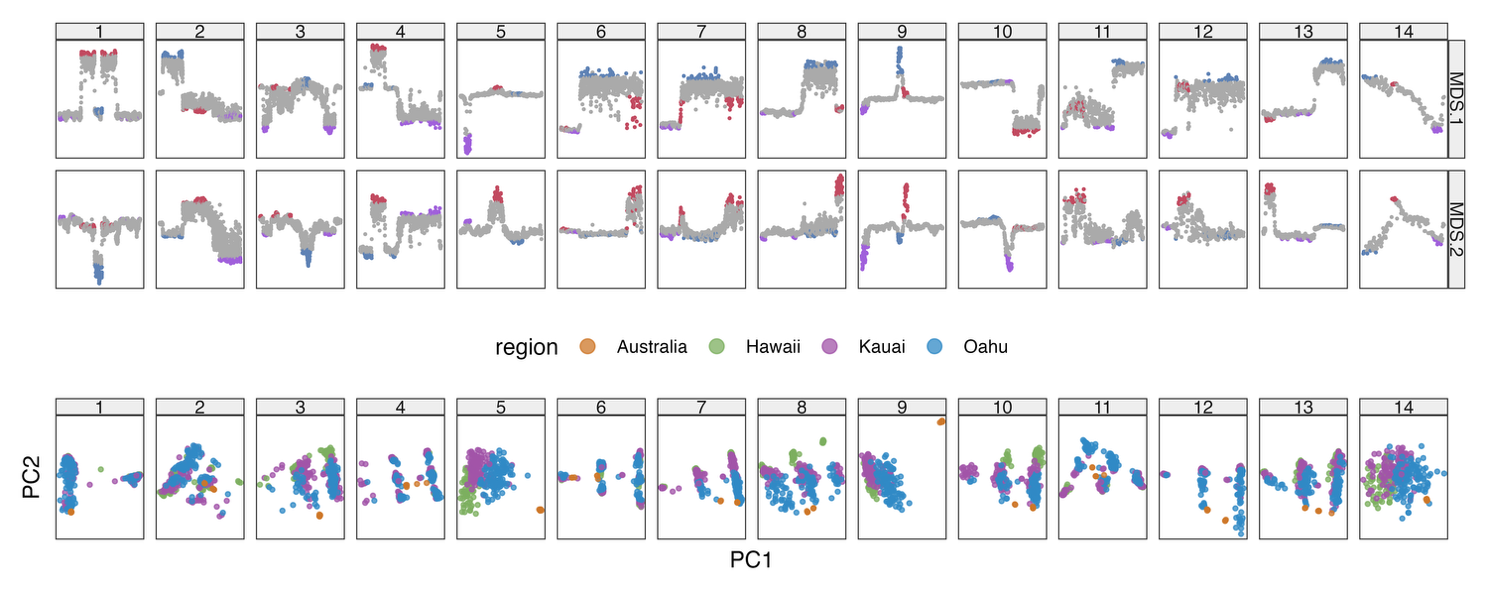

**Fig. S3.** Chromosome-wide PCA projections for samples from three Hawaiian islands, alongside seven further *T. oceanicus* samples from two mainland Australian populations. With the possible exception of the SV on chr6, we do not observe evidence of SVs identified in Hawaiian population in Australian samples.

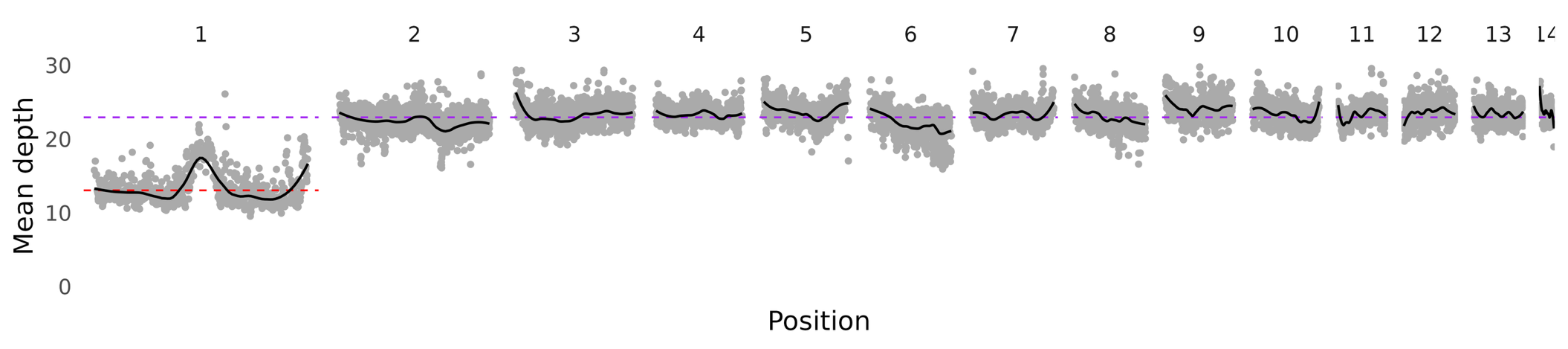
**Fig. S4.** Mean sequencing depth along each chromosome. Black lines show trends from LOESS regression. Purple lines show average read depth across filtered autosomal variants. The red line shows average read depth on Chr1, the X chromosome.

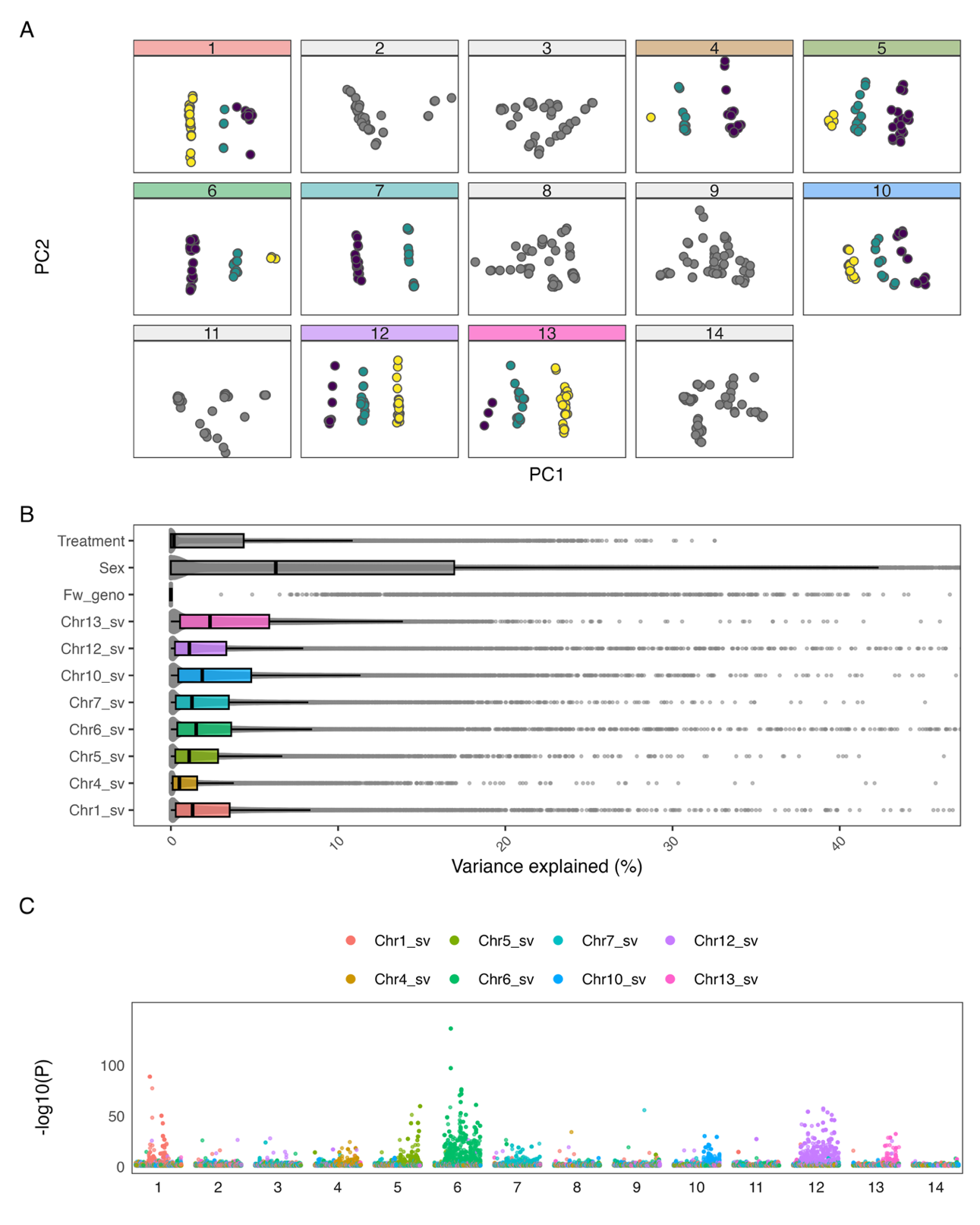

**Fig. S5. A)** PCA of filtered variants from RNA-seq data separated by chromosome, with discrete clustering of samples indicative of one or more large structural variants. Samples are coloured based on assigned karyotype. **B)** Distributions of the proportion of variation in gene expression across 25,359 genes that is accounted for by structural variants on different chromosomes (coloured), and other variables included in the original analysis by (*27*). *Treatment* refers to the presence of male cricket song playback during rearing (Y/N), and *Fw_geno* refers to genotype at the *flatwing* locus. **C)** The genomic distribution of genes included in the analysis, with greater values on the Y-axis indicative of stronger evidence of differential expression. Points are coloured according to the structural variant included as a predictor variable in the differential expression analysis. For example, the cluster of strongly differentially expressed green points on chromosome 6 indicate that the structural variant identified on this chromosome had strong effects on gene expression.

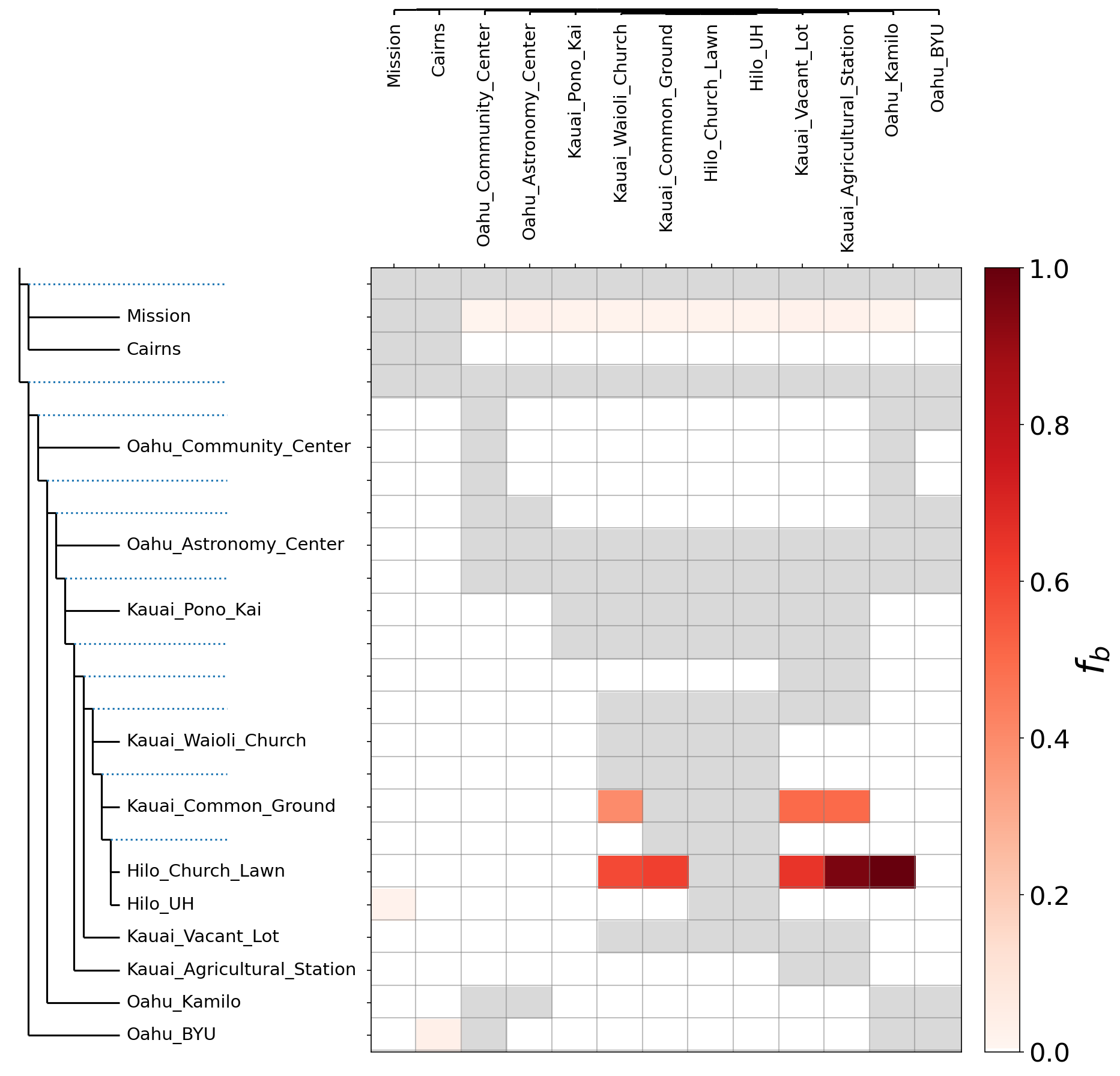

**Fig. S6.** F-branch statistics showing evidence of excess allele-sharing between multiple populations, indicative of gene flow.

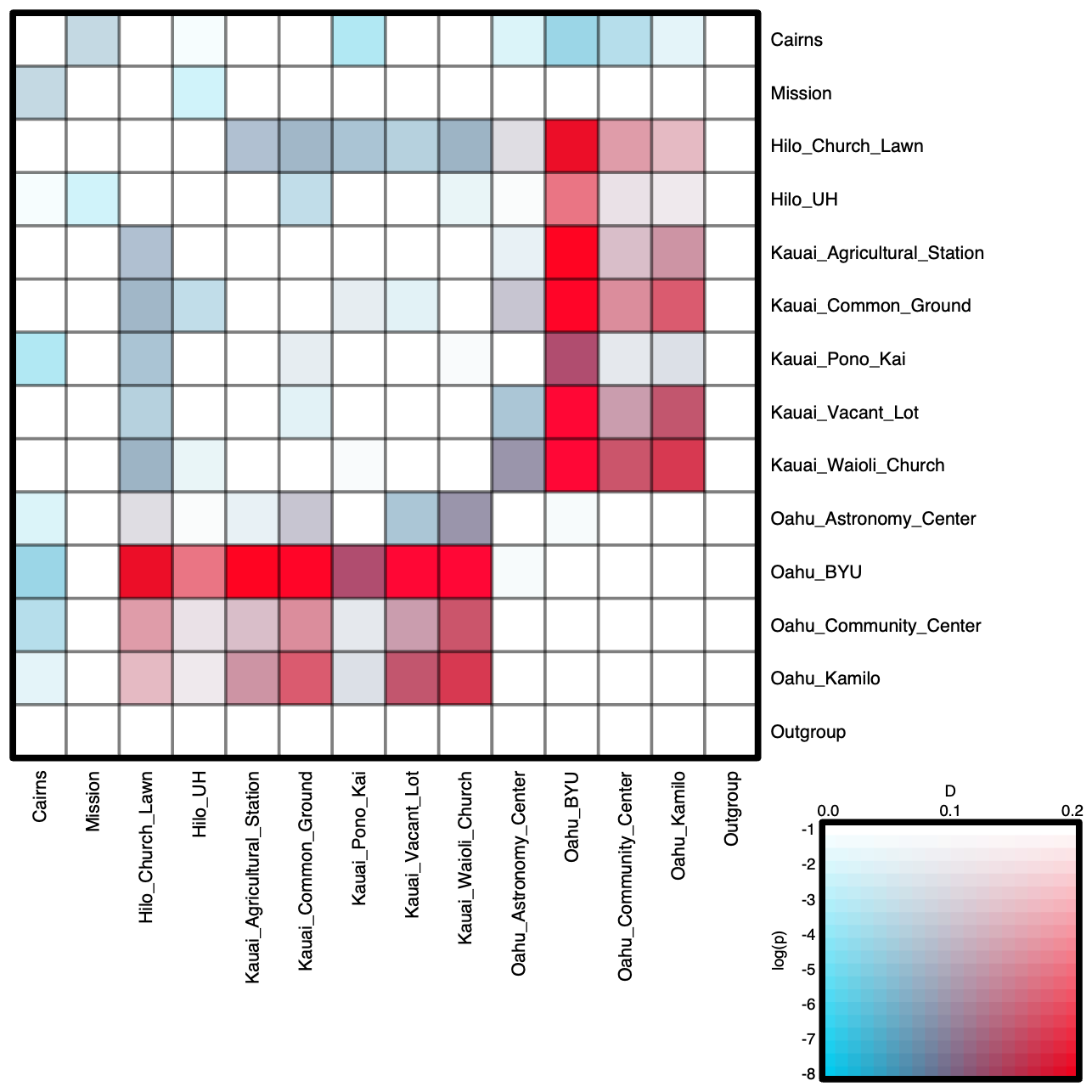

**Fig. S7.** ABBA-BABA statistics summarised across population pairs, coloured by D (hue) and P-value (saturation).

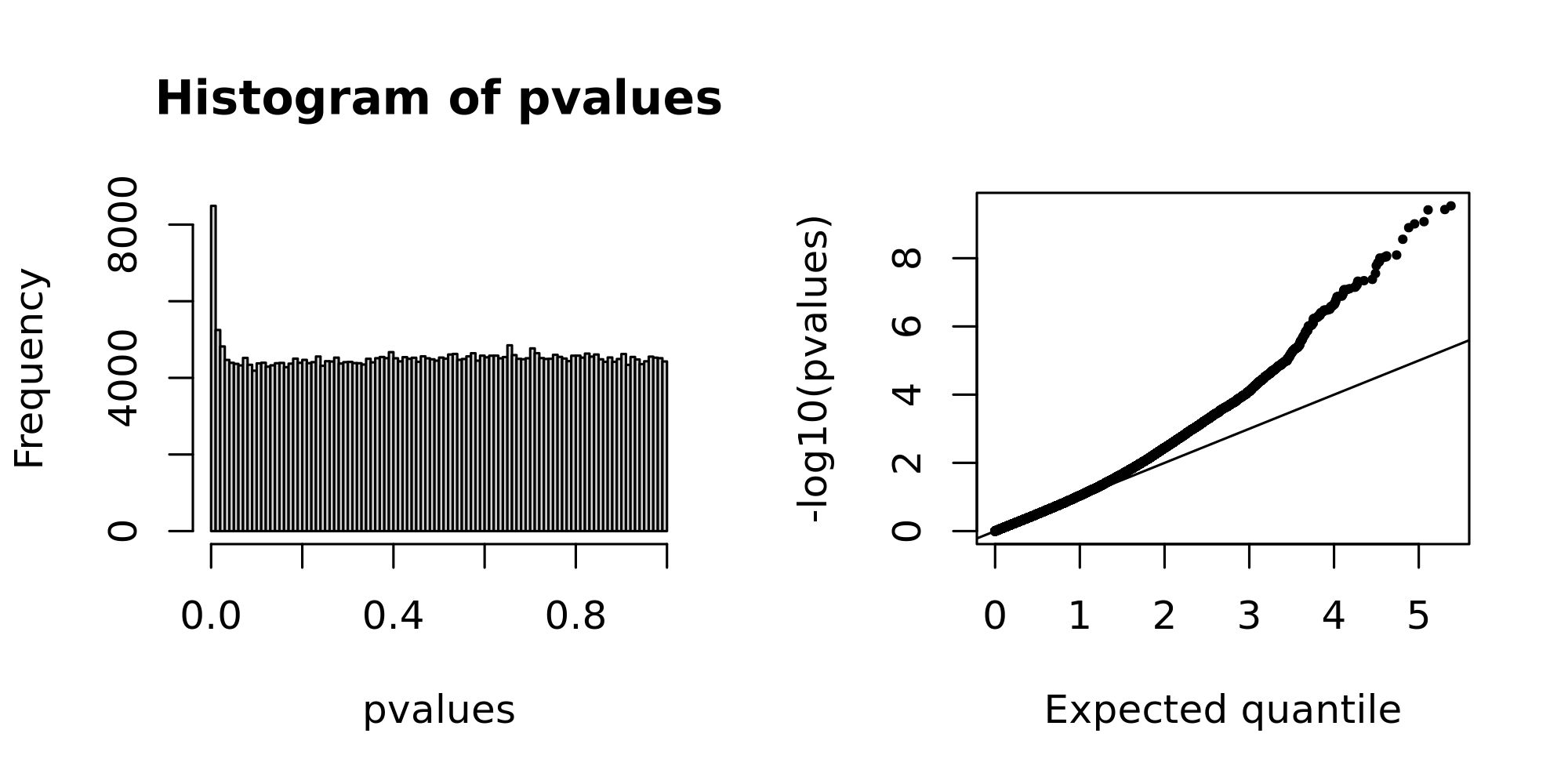

**Fig. S8.** A histogram of calibrated P-values from the LFMM used to test evidence of fly selection on cricket genome variants, and a qqplot illustrating the inflation of low P-values relative to expectation.
